# Unrelated Donor Selection for Stem Cell Transplants using Predictive Modelling

**DOI:** 10.1101/242735

**Authors:** Adarsh Sivasankaran, Eric Williams, Martin Maiers, Vladimir Cherkassky

## Abstract

Unrelated Donor selection for a Hematopoietic Stem Cell Transplant is a complex multi-stage process. Choosing the most suitable donor from a list of Human Leukocyte Antigen (HLA) matched donors can be challenging to even the most experienced physicians and search coordinators. The process involves experts sifting through potentially thousands of genetically compatible donors based on multiple factors. We propose a Machine Learning approach to donor selection based on historical searches performed and selections made for these searches. We describe the process of building a computational model to mimic the donor selection decision process and show benefits of using the proposed model in this study.

## I. Introduction

An Unrelated Donor (URD) search for Hematopoietic Stem Cell Transplant (HSCT) is initiated when patients are unable to find a match within their families. Studies have shown that 70% of the patients needing HSCTs have to rely on URDs or Cord-Blood Units (CBUs) (Gragert, et al. 2014). Volunteer donor registries and cord blood banks across the globe facilitate URD and CBU searches for such patients. Advances in HLA matching algorithms, therapeutic protocols, and large and diverse registries have resulted in increased number of HSCTs being preferred to treat patients (Copelan 2006). HLA compatibility between patient and URDs (and CBUs) is established by HLA matching algorithms (Bochtler et al. 2016). For URDs, a donor should match at least 8 of the 10 alleles at HLA-A, B, C, DQB1, DRB1 to be considered viable. A donor who match at all 10 alleles is the most preferred. After a list of suitable donors have been identified by matching algorithms, a physician or a search expert selects a short list of donors who are likely to provide the most optimal post-transplant outcome. This selection process is based on donors’ secondary characteristics (Spellman et al. 2012).

Optimizing clinically relevant criteria in the donor selection process for a stem cell transplant provides the best post-transplant outcomes. Depending on the HLA type of the patient the search process might involve selecting 3–5 donors for Confirmatory Typing (CT)^1^ from a long list of HLA matched URDs. Donor selection, subsequent to HLA matching, involves sifting through a long listing of URDs based on non-genetic factors.

Donor search and display systems, such as Traxis™ developed the National Marrow Donor Program (NMDP), are used to identify donors who are likely to provide the best post-transplant outcome. Donor characteristics that are identified by medical studies to provide favorable transplant outcomes are displayed in Traxis™ for the search experts to make an informed decision. A URD search for a patient with a common HLA, at NMDP, can potentially have tens of thousands of matched donors. Making a choice between identically matched donors can be an extremely long and difficult process and is done while the patient is under critical care. The selection process is based on evaluating multiple donor secondary characteristics (Spellman et al. 2012). The selection is also dependent on considerations of donor availability and Transplant Center (TC) experience with a Donor Center (DC), which cannot be measured and captured. A computational model that can help ease this selection process by quantitatively identifying donors with more preferable secondary characteristics based on past searches (Shouval et al. 2014). In this study, we develop a Machine Learning model that can mimic the donor selection process. Using a trained model, we can assign a *Selection Score* to every HLA matched donor for a patient to indicate favorability of secondary characteristics. Such a score will reduce the comparative multivariate decision process to a decision based on a single score that combines all the relevant donor features. It can be of particular assistance to physicians and TCs which lack the expertise and man-power to make such a critical decision (Irene et al. 2017).

Machine learning algorithms are being used in a variety of applications ranging from self-driving cars to financial forecasting. Computational models can be trained to mimic a specific human decision process, based on historical data without having to explicitly define and program the association between inputs and outputs. These trained models can then be used to predict the possible outcome of a future data instance. Figure 1a describe the decision process for donor selection and the modelling goal. In Figure 1b we show the proposed change to the process using a trained machine learning model. Based on historical searches and corresponding donor selections made we can train a computational model that imitates the donor selection process. The proposed model can make suggestions to the search experts to help in making the selection faster.

**Figure 1:**
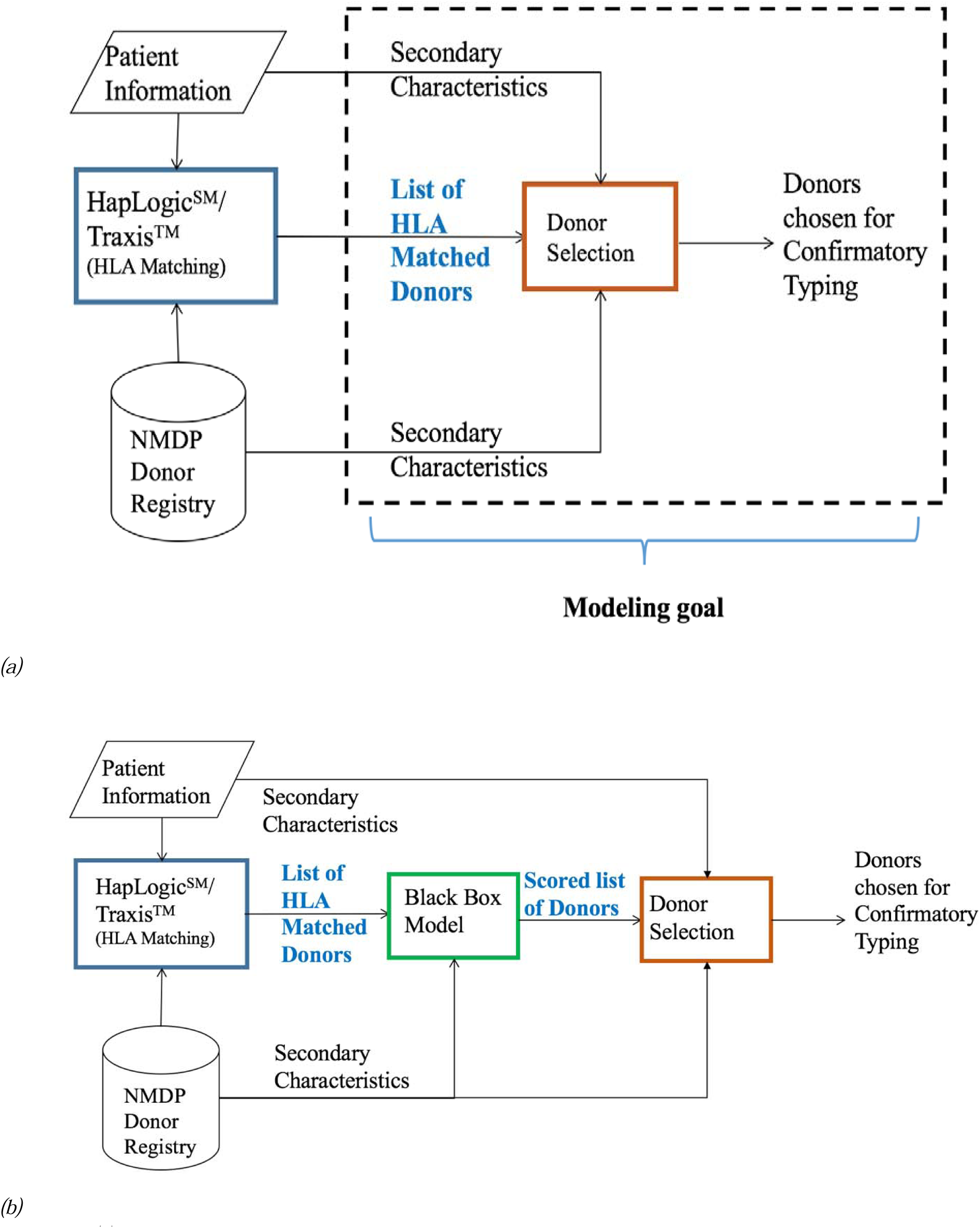
(a) Block diagram of the donor selection process. This study focuses entirely on modelling the selection decision process after HLA matching has been determined based on genetic population match probabilities. (b) This is the proposed system. The black box model will be used to score the list of HLA matched donors based on their secondary characteristics. The black box model can be integrated into the donor search system.

## II. Data and Methods

We have identified donor searches facilitated by the NMDP over a 3-month period (May 15, 2016 to Aug 9, 2016) for modelling. When a recipient and donor are fully marched (10/10 allele match), the considerations for donor selection are entirely based on non-genetic factors. For partially matched donors (8/10 and 9/10), number of mismatches and location of mismatch are factored into the selection. There are a number of studies that have separately identified effects of mismatched allele on the outcome of the transplant (Kanda et al. 2015) (Schetelig et al. 2008). A donor with more *suitable* mismatch locus is preferred over other donors (Nowak 2008). This translates to different selection criteria for fully matched and partially matched donors. Hence, these two scenarios need to be modelled separately. To facilitate this modelling aim, we identify all data that was presented in Traxis™ at the time a search was performed. This includes recipient-donor pair specific HLA matching information and donors’ non-genetic factors. HapLogic^SM^, the HLA matching algorithm developed by the NMDP, estimates an overall match probability as well as match probabilities at the individual allele level (Dehn et al. 2016). For partially matched donors at least 1 allele mismatch has been unambiguously identified. We collect the overall match probabilities (at 10/10, 9/10, and 8/10 match grades) and the mismatch locus information where applicable. This entire set of collected information is referred to as the secondary characteristics in this study. The ambiguity in donor HLA is resolved by high resolution typing during Confirmatory Typing.

In consultation with search experts at the NMDP we have identified donor characteristics that are important for the decision process. These are listed in Table 1 along with a description of any processing performed on these characteristics before modeling. Notice that several donor characteristics are missing in the database. This is due to either donors not sharing the complete information with NMDP when they were recruited or the information not being entered in the database.

**Table 1:** Donor Characteristics that are considered important to the donor selection process.

| Donor Characteristics | Significance | Considerations for Modelling |
| --- | --- | --- |
| *Donor's Age* | Younger donors are preferred over older donors (Kollman et al. 2016). | Information was available for 100% of the donors. Information was scaled on a [0,1] range before modelling |
| *Number of matched alleles* | At least 8 of the 10 alleles must be matched. 10/10 match is preferred over 9/10, which is preferred over 8/10 (Petersdorf et al. 2007). | This was only considered for modelling partially matched donor selections. A one-hot encoded variable was used to indicate match at 8,9, or 10 /10 match levels. |
| *Match Probability* | Assigned by HapLogic<sup>SM</sup> (Dehn et al. 2016) for every donor, in the range 1% - 99% for each level of match (i.e., 8/10, 9/10, or 10/10 level). A donor with higher match probability is preferred. Match probability at 10/10, 9/10 and 8/10 level are used appropriately. | HapLogic<sup>SM</sup> assigned match probabilities are retained as in. For fully matched modelling only 10/10 match probability is used. 8/10 and 9/10 match probabilities were considered for modelling partially matched donor selections. Variables were scaled on a [0,1] range. |
| *Donor's Blood type and Rh Factor* | When possible a match between recipient and donor's blood group and Rh factor is preferred (Worel 2016; Kollman et al. 2016). | This information was missing for about 24% of the donors in the dataset collected. Data values were altered to indicate if the information was present or absent at the time decision was made. Two variables were used to indicate Blood group and Rh type separately. |
| *Donor's Gender* | Male donors are favored (Spellman et al. 2012) | Encoded to a binary variable to indicate Male or Female. Complete information was available. |
| *Donor's Race* | Donors' Race is listed in Traxis<sup>TM</sup> (Dehn et al. 2016) as belonging to one of the following broad categories:<br>• African American (AFA)<br>• Asian/ Pacific Islander (API)<br>• Caucasian (CAU)<br>• Declined to Answer (DEC)<br>• Hispanics (HIS)<br>• Multi-Group (MLT)<br>• Native American Indian (NAM)<br>• Other Group (OTH)<br>• Unknown (UNK) | The categorical variable representing donor race was dummy encoded, with each category represented using an indicator variable. Groups DEC, MLT, OTH were merged with UNK category to account for data sparsity. Information was available for all donors. |
| *Donor's Weight* | This information is used to estimate the volume of stem cells that can be harvested from the donor, and if the donor can meet the requirements for the recipient. | This information was only present for 3.43% of the donors. We use a binary indicator to represent if the information was present or absent when the selection was made. |
| *Donor's CMV report* | CMV-seropositive or seronegative donors are preferred for CMV-seropositive and seronegative recipients, respectively | 11.97% donors had CMV screening results. We binary encode the information as |
|  | (Boeckh and Nichols 2004) | present or absent (binary). |
| *Low/high Resolution Typing* | High resolution typing indicates a stronger accuracy in match probabilities | Information was retained as a binary variable indicating high or low-resolution typing. |
| *DPB1 Permissivity* | A DPB1 permissive is preferred in addition to a 10-allele match. DPB1 match or a permissive mismatch is considered to be equally preferable (Fleischhauer et al. 2012; Shaw et al. 2014) | Information encoded in a binary variable to indicate match (permissive mismatch or a complete match) or not. |
| *Donor Chosen* | Information is collected to identify which of the matched donors is chosen to be asked for confirmatory typing. | Used as the target variable. A binary variable to indicate if a donor was chosen for the searched recipient or not. |

In addition to the variables listed in Table 1, we also collected Donor ID, Recipient ID, and Transplant Center ID for measuring model performance. These IDs are unique identifiers assigned by the NMDP for internal identification and communication. There are a few other factors, such as the previous pregnancy indicator for female donors, which did not have enough representation in the registry to be effectively modelled. Such factors have been ignored from consideration for this analysis.

For the specified 3-month period, NMDP facilitated a total of 2,138 donor searches. Among these (i) 1,439 searches had all their donor sections at the 10/10 level and (ii) 699 searches had at least 1 donor selected who was partially matched. For modeling case (i), we remove all partially (8/10, 9/10) matched donors since these were not considered during modelling. Donors who are at least 1% match at the 10/10 level are retained. In case (ii), we retain complete search results to model. These 2,138 donor searches resulted in a total of 8486 selections. That is an average of 4 selections per recipient. After data pruning, we have 1.5 million donor-recipient pairs for case (i) with 689k unique donors and 0.9 million donor-recipient pairs for case (ii) with 863k unique donors in the entire set. The difference in number of unique donors for the two cases is due to the heavy tailed distributions in HLA alleles as observed in (Slater et al. 2015). Two separate models are trained from these two sets of data. Table 2 has raw counts of chosen donors broken by key donor characteristics. This indicates selection preferences. For example, 61% of all chosen donors were male. We can also notice there is a definite preference for younger donors. 60% of all chosen donors were younger than 32 years of age, and only 7% of chosen donors were older than 50 years. Similarly, preferences can be inferred from Donor-Recipient HLA match data. Race is usually considered a difficult feature to determine (Hollenbach et al. 2015). Self-identified race information is often incorrect and do not correspond with genetic measurements. Machine Learning models do not require users to explicitly define these relationships.

**Table 2:** Raw count of chosen donors by various donor characteristics. This gives some indication of what the preference is. We see younger and male donors are preferred.

| Donor Broad Race Groups |  | Donor Gender |  |
| --- | --- | --- | --- |
| AFA | 499 | Female | 3279 |
| API | 534 | Male | 5207 |
| CAU | 4299 |  |  |
| DEC | 26 |  |  |
| HIS | 767 |  |  |
| MLT | 480 |  |  |
| NAM | 57 |  |  |
| OTH | 21 |  |  |
| UNK | 1803 |  |  |

| Donor Age |  |
| --- | --- |
| <= 32 | 5155 |
| [33-50] | 2717 |
| > 50 | 614 |

Donor-Recipient HLA Match Count
|  |  | Mismatched Location |  |  |  |  |  |
| --- | --- | --- | --- | --- | --- | --- | --- |
|  | Total | A | B | C | DQB1 | DRB1 |  |
| 10 on 10 | 6410 | - | - | - | - | - |  |
| 9 on 10 | 2000 | 1108 | 458 | 90 | 136 | 208 | $\Leftarrow 1^{st}$ Mismatch |
| 8 on 10 | 76 | 37 | 7 | 1 | 5 | 26 | $\Leftarrow 1^{st}$ Mismatch |
| | | 0 | 11 | 13 | 49 | 3 | $\Leftarrow 2^{nd}$ Mismatch |

Donor selection indicator is used to identify which donor was chosen to be the best match for a patient. That is, using the physicians’ choice indicator donors in the set were labelled as *chosen* or *not chosen* This labelling process allows us to formalize the learning problem under a binary classification setting (Cherkassky 2013). Binary classification is when data instances (donors, in our case) are to be classified into one of two groups (or *classes)*. Characteristics as listed in Table 1 are used to form a feature vector for each donor. A binary classifier is trained to identify a mathematical relation that will map donors to either of the two classes as above. A label *(chose* or *not chosen*) is then assigned to every donor based on their representative feature vector. Several models were trained and their accuracies were measured on an independent test set (out of sample data). SVMs were finally chosen based on their accuracy and computational run-time. For a thorough discussion on SVM please refer to (Burges 1998; Vapnik 2000; Cherkassky and Mulier 2007).

SVM is a very popular learning algorithm that has been used successfully for a variety of applications, including biomedical studies. (Furey et al. 2000) shows use of SVM for classifying cancer tissue samples from multiple microarray datasets. In (Y.-D. Zhang et al. 2015) SVM is used to predict recurrence of prostate cancer biochemical based on MR imaging and clinicopathology datasets. (H. Zhang et al. 2012) uses SVM to identify genes to improve cancer classification. (Verplancke et al. 2008) shows an application for SVM in hematological studies, where the goal is to predict mortality in critically ill patients. In comparative studies SVM performs as well or better than popular algorithms (Caruana and Niculescu-Mizil 2006).

For our problem, we also have a heavily *class-imbalanced dataset*. A dataset is imbalanced when number of samples in one class outnumber the number of instances in the other class. In our dataset, for every donor that is chosen about 300 donors remain not chosen. Modelling this huge discrepancy in classes poses challenges. Data classification techniques use a parameter called *misclassification costs* that inform the classifier how one type of error is penalized over the other, i.e., cost of making a false-positive versus cost of making a false-negative. These costs are completely dependent on the application and are supplied to the learning algorithm *apriori* (before modelling is performed). However, it is difficult to estimate the cost of making a mistake since we are dealing with patients’ care. We use ratio of classes as the misclassification cost. Data is separated into two sets: training set and testing set. Training data is used for model selection and test data is used for model evaluation. We use data from a total of 592 searches for model training. General experimental setup is as followed in (Sivasankaran et al. 2016), which has more detailed considerations of problem formalization, misclassification costs, multiple performance measurements, and model selection criteria used in this modelling effort. A number of publicly available packages have SVM implementation. An R Package called ‘LiblineaR’ (Fan et al. 2008) was used. Further analysis shown in the following section is only based on out-of-sample data (test data) from 1546 searches.

## III. Results and Discussion

SVM classifiers are typically used to assign a predicted label to new data instances. Use of hard label assignment will lead to donors being labelled *chosen* or *not chosen*. The end goal of this modelling is to be able to suggest donors most likely to be asked to donate with a higher score. Hard label assignment will not help us with this goal. SVMs use something called projection distance to make the hard label assignments. These distances are assigned based on the distance between a data instance and the learned separating hyperplane as shown in Figure 2. This distance can be used to assign a real valued score to matched donors instead of class labels.

**Figure 2:**
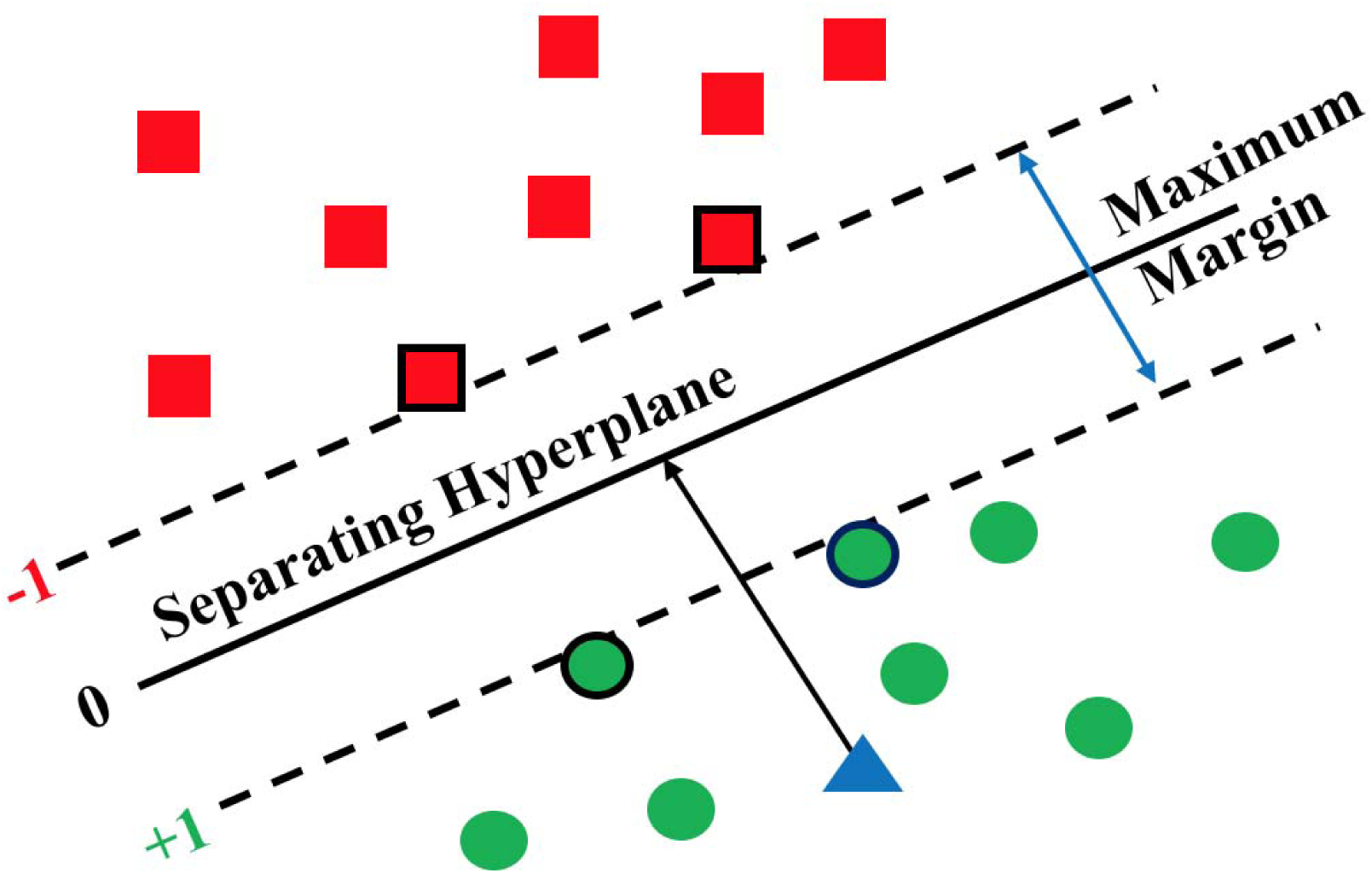
An illustrative example of a SVM hyperplane. Green circles form the +1 class and the red squares belong to −1 class. The markers on each side with blackened boundaries are called Support Vectors. The blue triangle is a new point that needs a class assignment. Based on its location with respect to the plane, it will be assigned to the +1 class. The margins on either side will be equidistant from the separating hyperplane.

To effectively assist the decision process, donors who are more likely to be chosen should be assigned a higher score than other donors in the list, i.e., donors with favorable secondary characteristics should receive higher scores than the donors with less favorable characteristics. The proposed SVM model is trained on past donor selections. Figure 3 shows the described sorting method for a hypothetical donor search. A list of HLA compatible donors as identified by the matching algorithm are assigned a score based on their secondary characteristics. This score is then used to sort the donors for the donor display system. This may help TCs to make decisions faster by quantitatively defining favorability of donors’ secondary characteristics.

**Figure 3:**
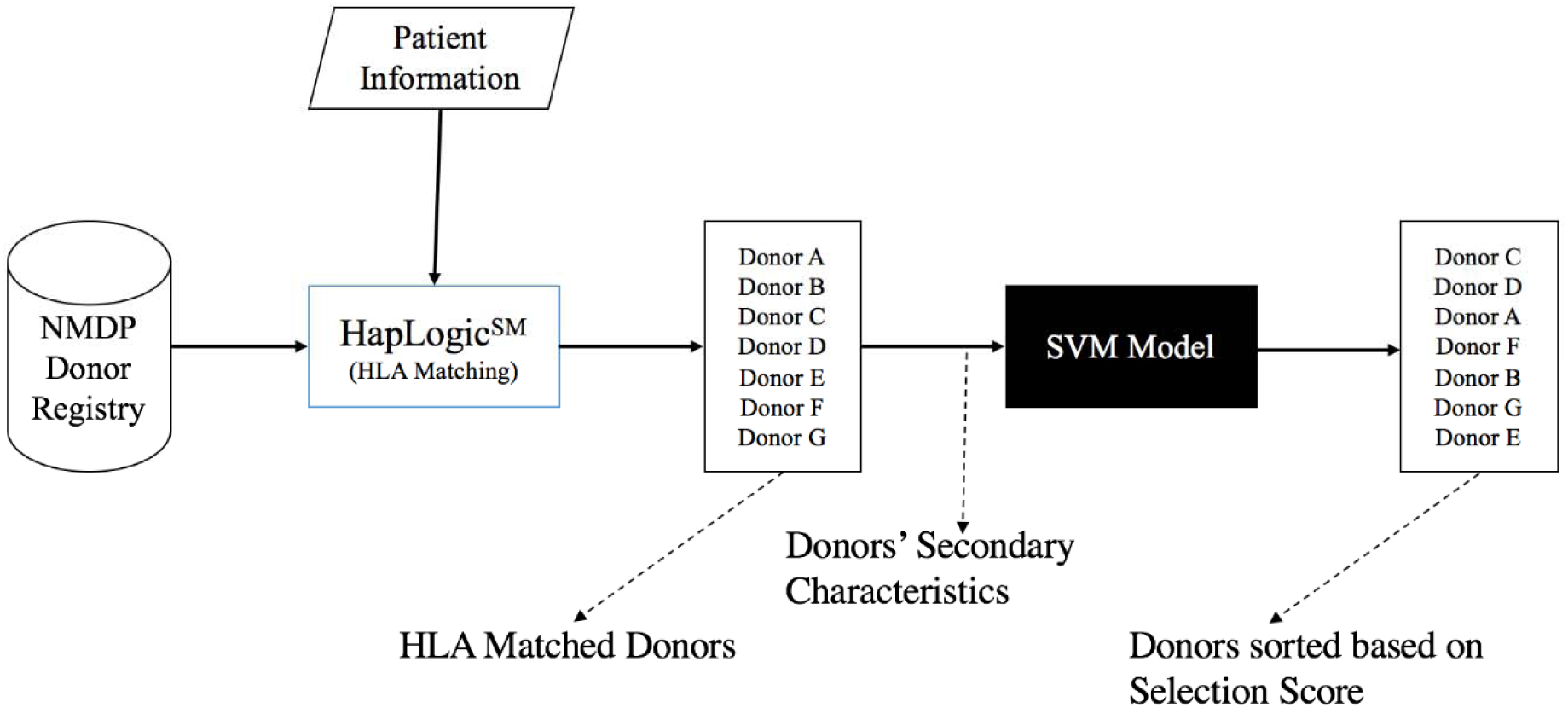
Donor sorting based on SVM model. Once the donors are identified to be a match for a patient's HLA type from the registry, the donor's secondary characteristics are fed to the trained SVM model to obtain a score for each donor. The donors are then sorted in ascending order based on the score.

*Search Presentation:* Using SVM projection distances, a graphical representation of the high dimensional data can be produced using a novel technique called Histogram of Projections (Cherkassky 2013) for each patient as shown in Figure 4. This representation provides the users with an ability to view the multi-variate high dimensional donor data on a simple histogram. This will help hugely narrow down the number of donors that need to be considered to make the decision. Figure 4a and 4b show the histograms of projections for real patients with 415 and 456 matched donors respectively, that were not used for training the model. Using the histogram of projections, can restrict the search size from the entire list to a handful of donors with positive scores. In Figure 4a, a small portion of the donors have really high scores, indicating donors with really favorable characteristics. In Figure 4b, donor scores are clustered, indicating donors have similar secondary characteristics. Here too, search field can be limited only to donors with positive scores. This representation can be easily integrated to the donor display system. Donors with scores higher than +1 will have extremely favorable characteristics, and similarly, and donors with scores less than −1 will have unfavorable characteristics. We notice that in most searches only a small percentage of matched donors were assigned positive scores.

**Figure 4:**
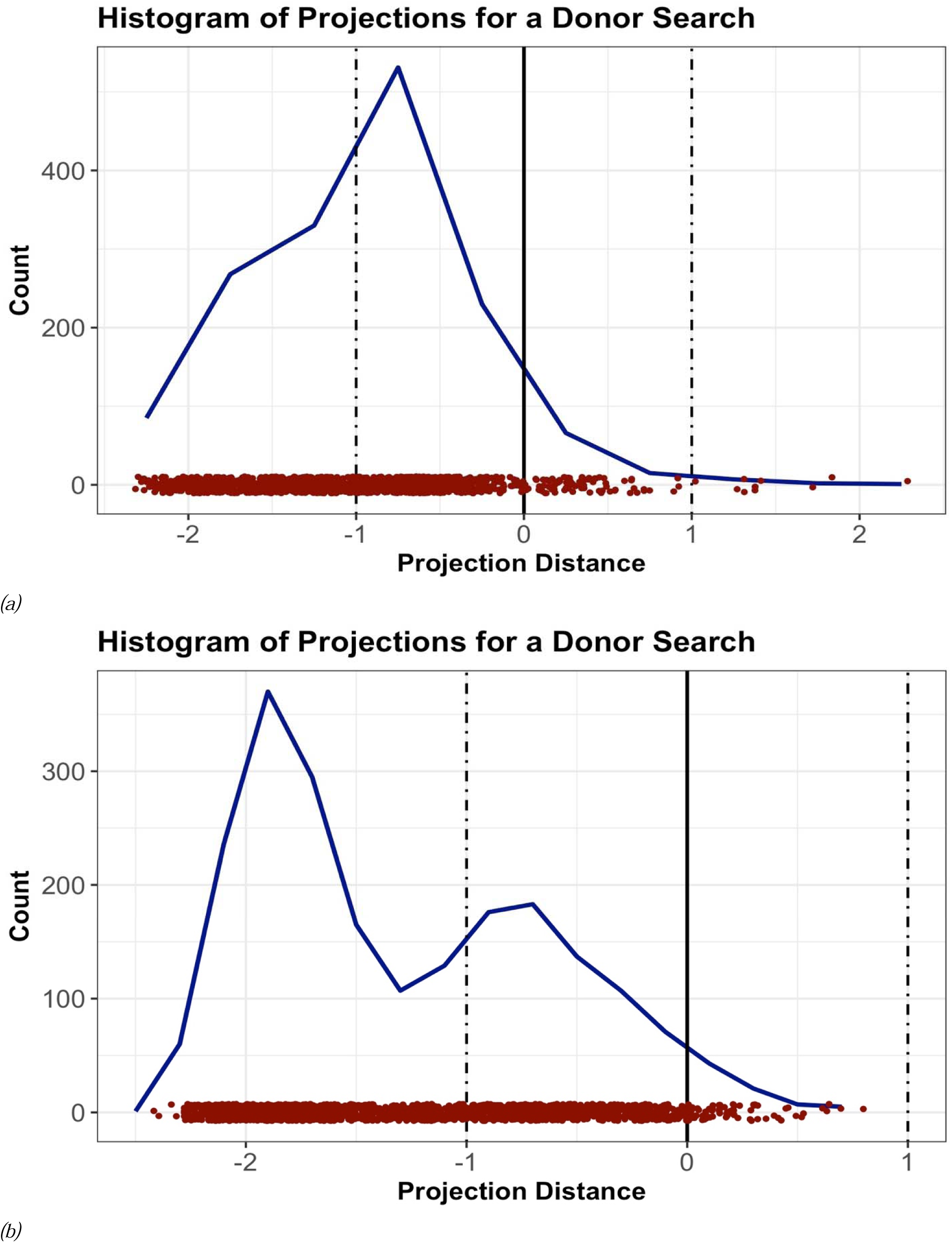
(a) Histogram of Projections for a search with 415 matched donors. A small portion of donors have a very high score, (b) histogram of Projections for a search with 456 donors, with a more clustered score.

After data cleaning as described in Section 2, the maximum number of matched donors for a search was 44,646. Looking for the best donor for this patient would have been extremely time consuming. Donor search experience can be improved using the modelled score. Figure 3 shows how SVM model can be used to sort donors. Matched donors are assigned a rank based on decreasing model score – donor with the highest score gets rank 1, donor with the second highest score gets rank 2, and so on. We analyze ranks of chosen donors (per patient) based on the proposed sorting method. Figure 5 has the cumulative distribution of the maximum rank (position of donors in the sorted list) of chosen donors. 75% of all the searches had all their chosen donors ranked within a position of 45.

**Figure 5:**
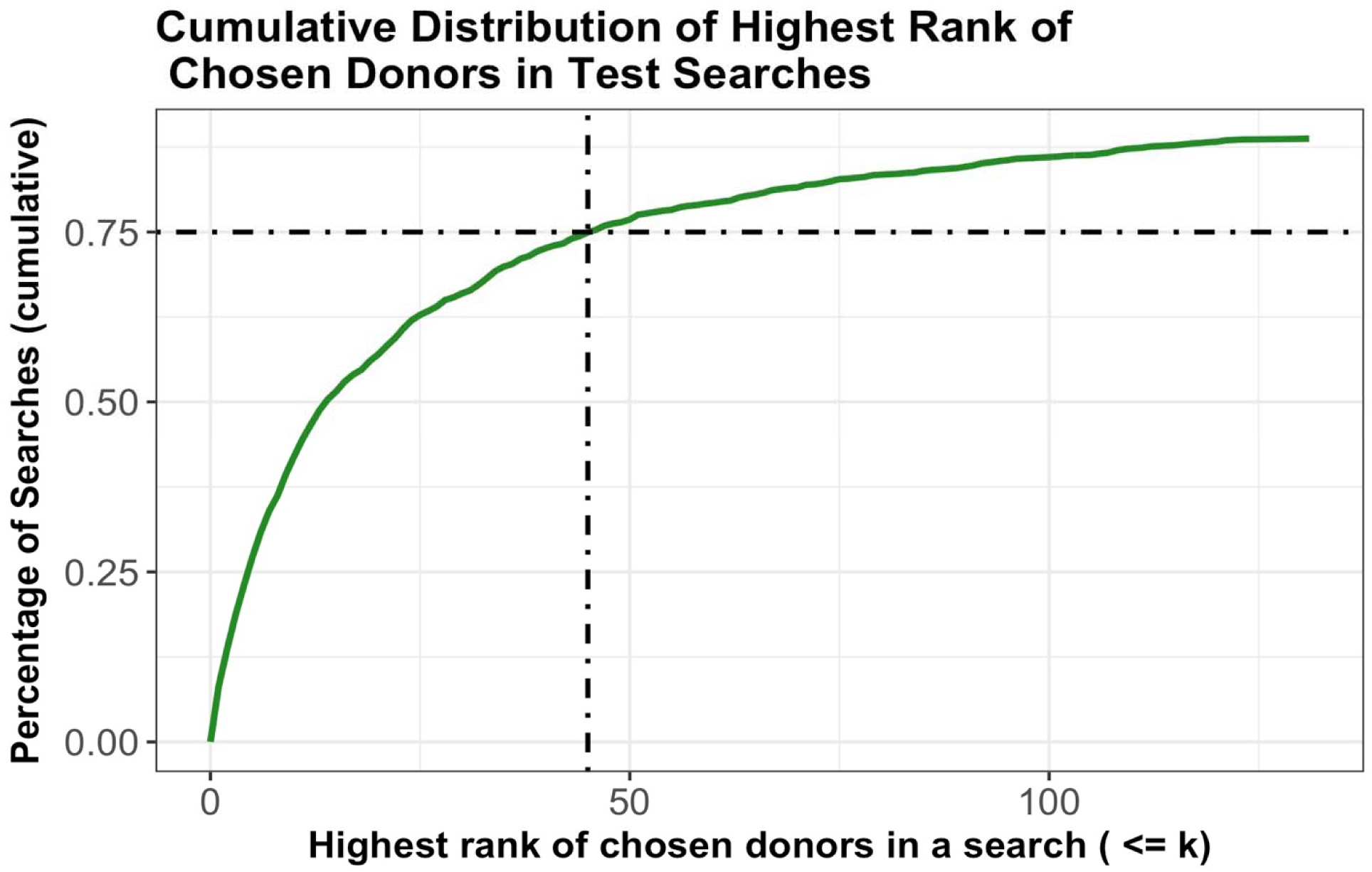
Cumulative distribution of highest rank of chosen donors. The graph is truncated on the x-axis for presentation.

*Transplant Center Monitoring:* Donor selection requires dedicated staff with expertise and knowledge of selection protocols and is a time-consuming process. The current modelling effort provides a direct method to quantify the search efficiency. Model assigned scores can be used to analyze donor selection behavior of TCs and provide feedback when it is noticed suboptimal choices are made repeatedly. Figure 6 shows a hypothetical behaviour for two Transplant Centers. An optimal selection is when most of TC’s selected donors have a positive selection score. Not all selected donors will have a positive selection score. This happens when a choice has to made between donors with equally unfavorable secondary characteristics. Inconsistent donor selection practice will also lead to donors with unfavorable secondary characteristics being chosen over donors with more favorable characteristics. A similar density plot can be estimated for donor selections made by each TC and analyze selection behaviour.

**Figure 6:**
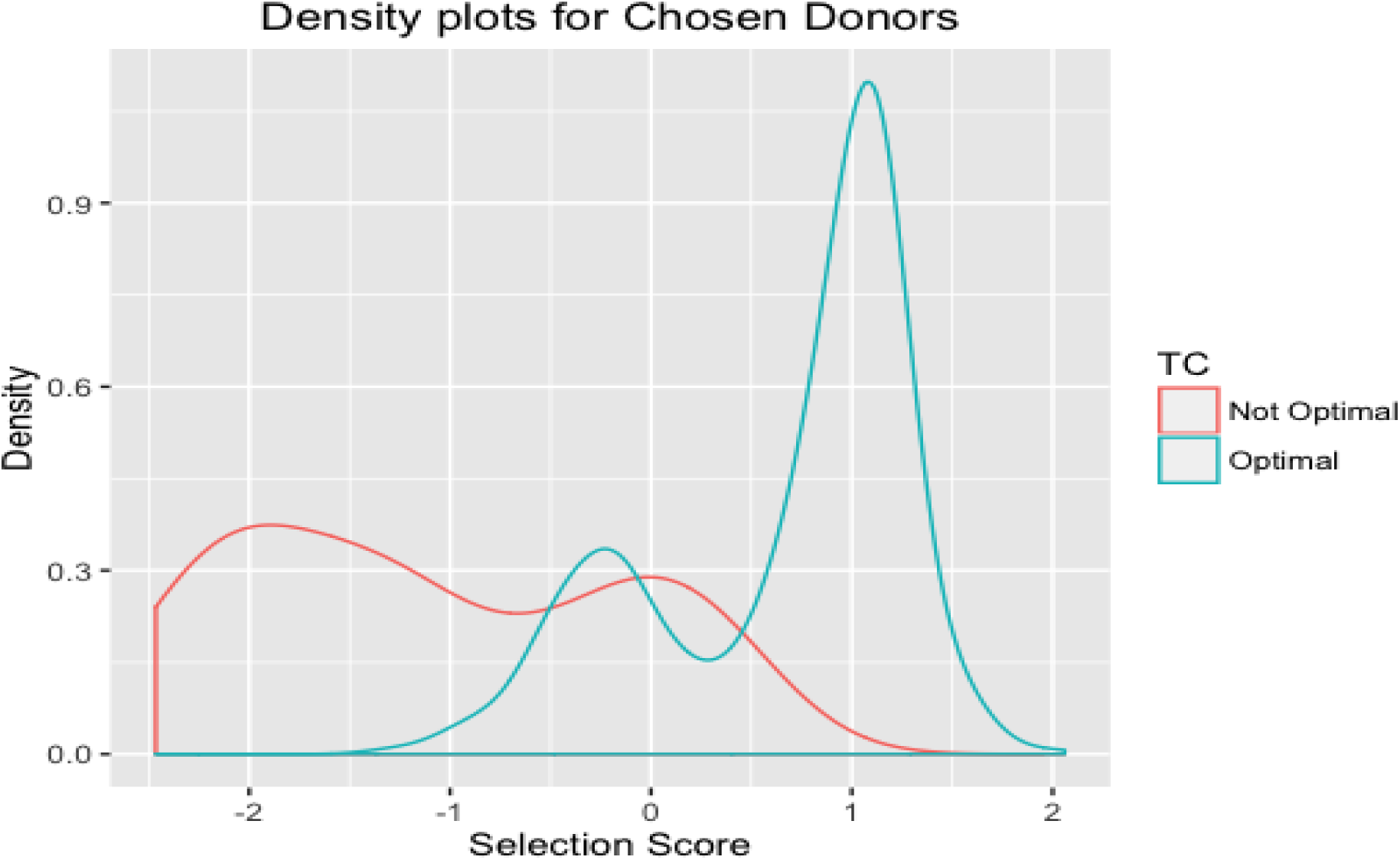
Selection Score densities for Optimal and Not Optimal choices

## IV. Conclusion

Donor searches are often performed when patients are under critical care. Having to choose between identically matched donors can be a huge burden on physicians and search experts. We have shown that use of Machine Learning can alleviate some of this pain and help make decisions faster. The trained model provides a quantitative way to compare and choose donors and the decision can be reduced to a single variable. This will help in making choices faster and complete transplants quickly. Further analysis of variable weights has shown that they correspond to how decisions are made in practice. Incorporating the model information into donor display systems can help streamline the URD search process and improve efficiencies. Time to transplant is an important concern for both TCs and Donor Registries (Dehn et al. 2016). The proposed model promises to reduce time spent on reviewing search results to make the most suitable choice.

## Footnotes

1 Members on the registry have ambiguous typing information. A probability is assigned to possible genotypes a member can potentially have. Confirmatory Typing is performed to resolve the ambiguity and confirm the member’s genotype.

